# CD8^+^ T cell signature in acute SARS-CoV-2 infection identifies memory precursors

**DOI:** 10.1101/2021.07.22.453029

**Authors:** Sarah Adamo, Jan Michler, Yves Zurbuchen, Carlo Cervia, Patrick Taeschler, Miro E. Raeber, Simona Baghai Sain, Jakob Nilsson, Andreas E. Moor, Onur Boyman

## Abstract

Immunological memory is a hallmark of adaptive immunity and facilitates an accelerated and enhanced immune response upon re-infection with the same pathogen^1,2^. Since the outbreak of the ongoing coronavirus disease 19 (COVID-19) pandemic, a key question has focused on whether severe acute respiratory syndrome coronavirus 2 (SARS-CoV-2)-specific T cells stimulated during acute infection give rise to long-lived memory T cells^3^. Using spectral flow cytometry combined with cellular indexing of transcriptomes and T cell receptor (TCR) sequencing we longitudinally characterize individual SARS-CoV-2-specific CD8^+^ T cells of COVID-19 patients from acute infection to one year into recovery and find a distinct signature identifying long-lived memory CD8^+^ T cells. SARS-CoV-2-specific memory CD8^+^ T cells persisting one year after acute infection re-express CD45RA and interleukin-7 receptor α (CD127), upregulate T cell factor-1 (TCF1), and maintain low CCR7, thus resembling CD45RA^+^ effector-memory T (T_EMRA_) cells. Tracking individual clones of SARS-CoV-2-specific CD8^+^ T cells, we reveal that an interferon signature marks clones giving rise to long-lived cells, whereas prolonged proliferation and mammalian target of rapamycin (mTOR) signaling are associated with clone contraction and disappearance. Collectively, we identify a transcriptional signature differentiating short-from long-lived memory CD8^+^ T cells following an acute virus infection in humans.

---

The COVID-19 pandemic has taken an unprecedented toll on global health and economy, impacting billions of lives all over the world. Importantly, the ongoing vaccination efforts appear to curtail the spread of SARS-CoV-2 and prevent severe disease, even as new virus variants emerge^4,5^. Yet, a prevailing question concerns whether and how exposure to SARS-CoV-2 by infection or immunization might result in long-term protective immunity.

The immune system reacts to an acute virus infection by mounting a rapid innate immune response, which is typically followed by stimulation of adaptive immune cells, including T cells and antibody-producing B cells. On encountering their cognate antigen on antigen-presenting cells, antigen-specific T cells vigorously proliferate and differentiate into effector cells aimed at controlling the pathogen and killing virus-infected host cells. Following elimination of the virus, 90-95% of effector T cells undergo apoptosis, whereas some antigen-specific T cells survive to become long-lived memory T cells able to protect the host from re-infection with the same pathogen^2^. While antigen-specific effector T cell responses are generated during acute SARS-CoV-2 infection^6–8^ and persist for several months^9–11^, little is known about the factors instructing individual effector T cell clones on their development to long-lived memory T cells in humans following an acute viral infection, as most of our current knowledge stems from animal studies^12^, *in vitro* experiments using human T cells^13^, or humans following vaccination^14,15^.

To determine which CD8^+^ T cell features present during acute infection are predictive of acquisition of memory and persistence several months after infection, we profiled SARS-CoV-2-specific CD8^+^ T cells directly *ex vivo* by using spectral flow cytometry, single cell RNA sequencing (scRNAseq) and TCR sequencing during acute COVID-19 as well as six and 12 months after primary infection.

## Phenotype of SARS-CoV-2-specific CD8^+^ T cells in acute COVID-19

To assess the dynamics of antigen-specific T cells in COVID-19, we recruited 175 patients with real-time polymerase chain reaction (RT-PCR)-confirmed COVID-19, sampled during their symptomatic acute phase and followed up six months and one year after acute infection (Fig. 1a). We conducted human leukocyte antigen (HLA) typing on all patients and healthy controls and selected individuals carrying the HLA-A*01:01, HLA-A*11:01 or HLA-A*24:02 alleles for this study (*n* = 33 patients, *n* = 13 healthy controls). In these individuals, SARS-CoV-2-specific CD8^+^ T cells were detected by using HLA-A*01:01, HLA-A*11:01 and HLA-A*24:02 major histocompatibility complex class I (MHC-I) dextramers^16^, hereafter collectively termed CoV2-Dex (Fig. 1b and Extended Data Fig. 1a), and validated by using HLA-A*01:01 and HLA-A*11:01 MHC-I pentamers^17^, hereafter termed CoV2-Pent (Extended Data Fig. 1b,c). SARS-CoV-2-specific CD8^+^ T cells were found in most patients in the acute phase and six months after acute infection (Fig. 1c), confirming previous findings using SARS-CoV-2 peptide stimulation^9–11,18–22^. Moreover, we detected SARS-CoV-2-specific CD8^+^ T cells one year after acute SARS-CoV-2 infection (Fig. 1b,c). As observed for murine virus infection^23^, the frequency of CoV2-Dex^+^ cells in the acute infection correlated with the frequency of specific cells during the memory phase (Fig. 1d).

**Figure 1.**
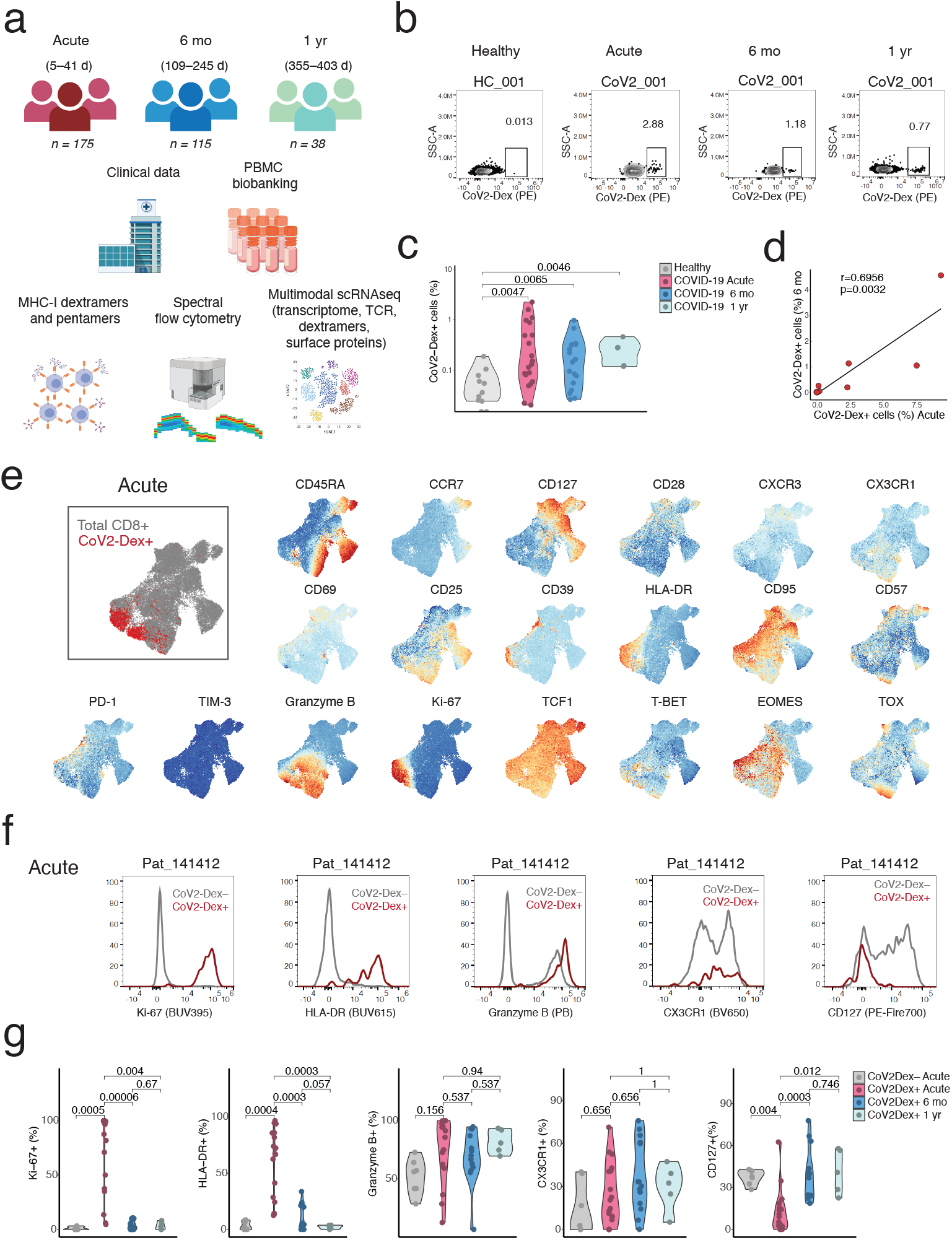
Characterization of antigen-specific CD8^+^ T cells during acute and memory phases of SARS-CoV-2 infection. **a**, Overview of study design. **b**, Representative plots of SARS-CoV-2 directed MHC-I dextramers (CoV2-Dex) staining. **c**, Percentage of CoV2-Dex^+^ cells in healthy donors and COVID-19 patients in the acute phase as well as six months and one year after infection. Each dot represents independent and unrelated donors (*n* = 24 acute, *n =* 22 six months after infection, *n =* 5 one year after infection), samples stained with HLA-A*24:02 dextramers were excluded due to a strong background staining on cells from unexposed individuals. **d**, Linear regression of frequency of CoV2-Dex+ cells six months after primary infection as a function of frequency of CoV2-Dex+ cells during the acute phase (patients sampled before day 14 since onset of symptoms were excluded from this analysis). **e**, Uniform manifold approximation and projection (UMAP) plots of marker expression for up to 2,000 CD8^+^ T cells from each sample analyzed by spectral flow cytometry. Regions with high expression of specific markers appear red. Overlay of CoV2-Dex^+^ cells (red) and Total CD8^+^ T cells (gray) is shown on the upper left. **f**, Representative histograms showing expression of selected markers on CoV2-Dex^−^ and CoV2-Dex^+^ cells. **g**, Frequency of Ki-67^+^, HLA-DR^+^, Granzyme B^+^, CX3CR1^+^ and CD127^+^ cells among CoV2-Dex^−^ (grey) and CoV2-Dex+ during acute infection, six months and 1 year after infection (red, blue and cyan, respectively). **c**, P-values were calculated with a Mann-Whitney-Wilcoxon test. **g**, P-values were calculated with a Mann-Whitney-Wilcoxon test and corrected for multiple comparisons.

In the acute phase, flow cytometry analysis of CoV2-Dex^+^ cells revealed a circumscript phenotype of activated cells, dominated by very high abundance of Ki-67 and HLA-DR (Fig. 1e and Extended Data Fig. 1e,f). CoV2-Dex^+^ cells also tended to express granzyme B and the terminal differentiation marker CX3CR1, whereas surface CD127 was markedly downregulated on CoV2-Dex^+^ cells (Fig. 1f). At the six-month and one-year timepoints, the frequencies of Ki-67^+^ and HLA-DR^+^ CoV2-Dex^+^ cells declined and the frequency of CD127^+^ cells increased, indicating a transition from effector to memory state, while granyzme B and CX3CR1 levels remained stable (Fig. 1g, Extended Data Fig. 2).

To examine the transcriptional phenotype of SARS-CoV-2-specific CD8^+^ T cells, we sorted CoV2-Dex^+^ CD8^+^ T cells and CoV2-Dex^−^ CD8^+^ T cells, mixed them at a 1:10 ratio, and performed scRNAseq on a subgroup of patients (*n* = 20 acute, *n* = 19 six-month timepoint). We classified sequenced cells as CoV2-Dex^−^ or CoV2-Dex^+^ based on their dCODE dextramer unique molecular identifier (UMI) counts (see Methods section and Extended Data Fig. 3a) and positivity for a single SARS-CoV2 epitope (Extended Data Fig. 3b). Unbiased clustering revealed 12 distinct CD8^+^ T cell clusters (Fig. 2a), none of which was dominated by a single patient (Extended Data Fig.4). Some clusters showed nearly complete segregation between acute and memory phase. Thus, cluster 12 was almost exclusive to acute COVID-19, whereas cluster 11 was mainly detectable in the convalescent patients (Fig. 2b,c). In line with our flow cytometry data (Fig. 1e and Extended Data Fig. 5), CoV2-Dex^+^ CD8^+^ T cells showed a more defined transcriptional makeup in the acute phase, whereas their transcriptional expression was more heterogeneous six months after infection (Fig. 2d). Comparing the contribution of CoV2-Dex^+^ cells to different clusters, we observed that clusters 1, 2 and 12 dominated the CoV2-Dex^+^ CD8^+^ T cell response in the acute phase, whereas cluster 11 contained CoV2-Dex^+^ cells exclusively from the recovery phase (Fig. 2e and Extended Data Fig. 6). While clusters 1, 2 and 12 corresponded to cytotoxic, activated, and proliferating cells, respectively, cluster 11 showed a dual signature marked by enrichment of interferon (IFN) response genes and genes encoding the effector cytokines IFN-γ, tumor necrosis factor (TNF), and lymphotoxin-α (LT-α) (Fig. 2f,g). Collectively, these data show that SARS-CoV-2-specific CD8^+^ T cells are predominantly activated and proliferating cells with a cytotoxic phenotype during acute COVID-19. However, at the six-month and one-year timepoints after infection, activation and proliferation subside and other hallmarks typical of resting memory T cells emerge, such as CD127 expression^24,25^.

**Figure 2.**
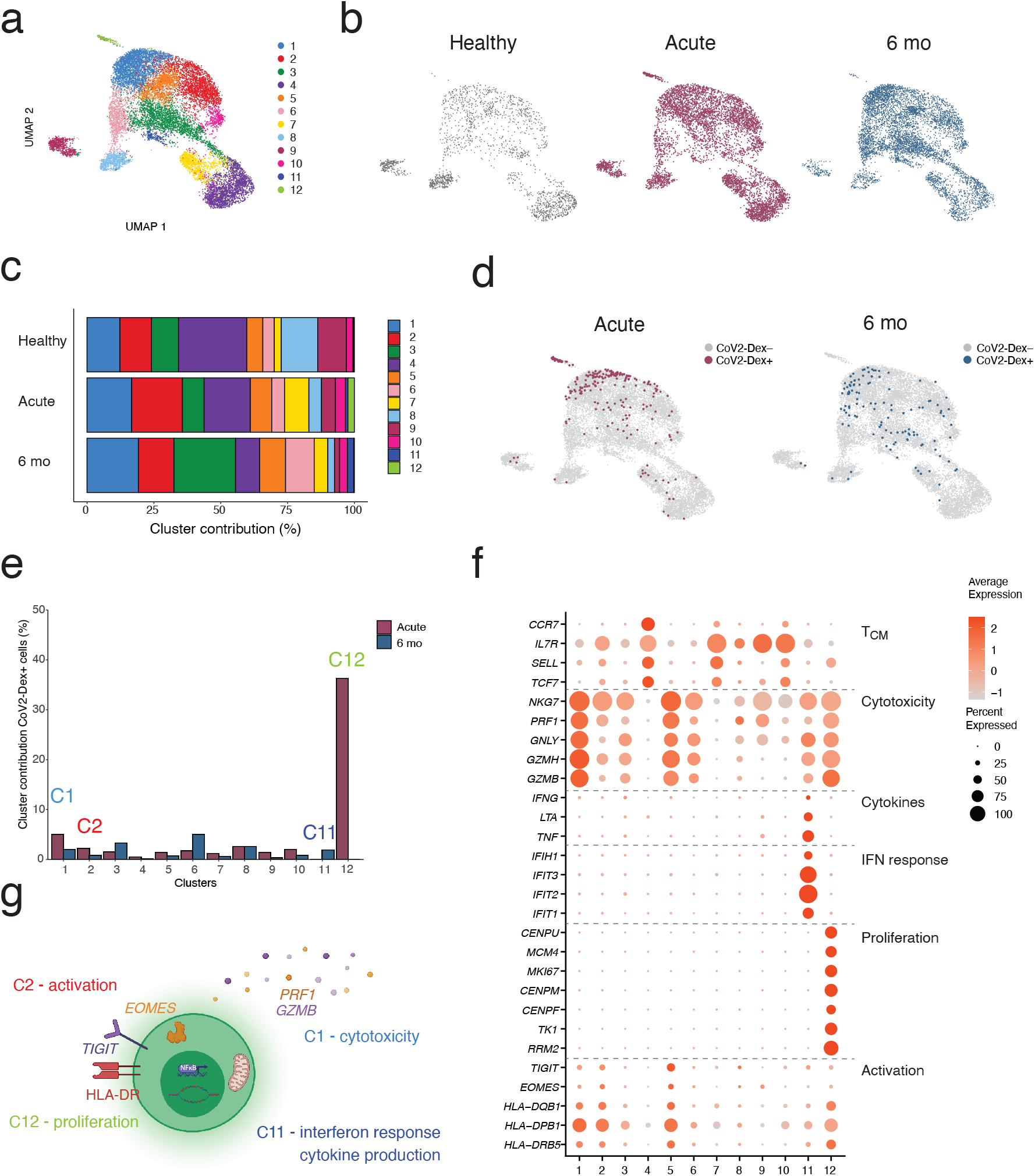
Transcriptional makeup of SARS-CoV-2-specific CD8^+^ T cells. **a**, Single-cell transcriptomes of CD8^+^ T cells displayed by UMAP. Seurat-based clustering of 14853 cells colored based on cluster ID. **b**, UMAP plot of CD8^+^ T cells of our study colored by status (healthy, acute COVID-19 or convalescent six months after infection). **c**, Cluster contribution to total CD8^+^ T cells of healthy, acute COVID-19 patients or convalescent patients six months after infection. **d**, UMAP of CoV2-Dex^−^ and CoV2-Dex^+^ single-cell transcriptomes in the acute phase and six months after primary infection. **e**, Percentage of CoV2-Dex^+^ cells contributing to each cluster either in the acute phase or six months after primary infection (percentages are calculated on total cells per cluster per timepoint). **f**, Average expression (color scale) and percent of expressing cells (size scale) of selected genes in each cluster. **g**, Schematic summary of the main clusters differentially represented in the acute phase and six months after primary infection.

## Phenotypic memory trajectories following SARS-CoV-2 infection

While a few seminal studies have explored phenotypic differentiation trajectories of antigen-specific CD8^+^ T cells in vaccinated human subjects^14,15^, little is known about the temporal changes of longitudinally tracked human antigen-specific CD8^+^ T cells following an acute infection. Thus, we aimed at identifying phenotypic differentiation trajectories of SARS-CoV-2-specific CD8^+^ T cells, since our longitudinal sampling allowed such analysis. In the acute phase, CoV2-Dex^+^ cells showed mostly an effector/effector-memory (T_effector_/T_EM_) phenotype, whereas frequencies of naïve (T_naïve_) and central memory (T_CM_) cells were lower in CoV2-Dex^+^ compared to CoV2-Dex^−^ CD8^+^ T cells (Fig. 3a,b, Extended Data Fig. 7a). These data were confirmed in CoV2-Pent^+^ cells (Extended Data Fig. 7b,c). At the six-month and one-year timepoints after infection, we observed a progressive switch from a T_effector_/T_EM_ to a T_EMRA_ phenotype, thus one year after acute COVID-19, most CoV2Dex^+^ cells were of a T_EMRA_ phenotype (Fig. 3c).

**Figure 3.**
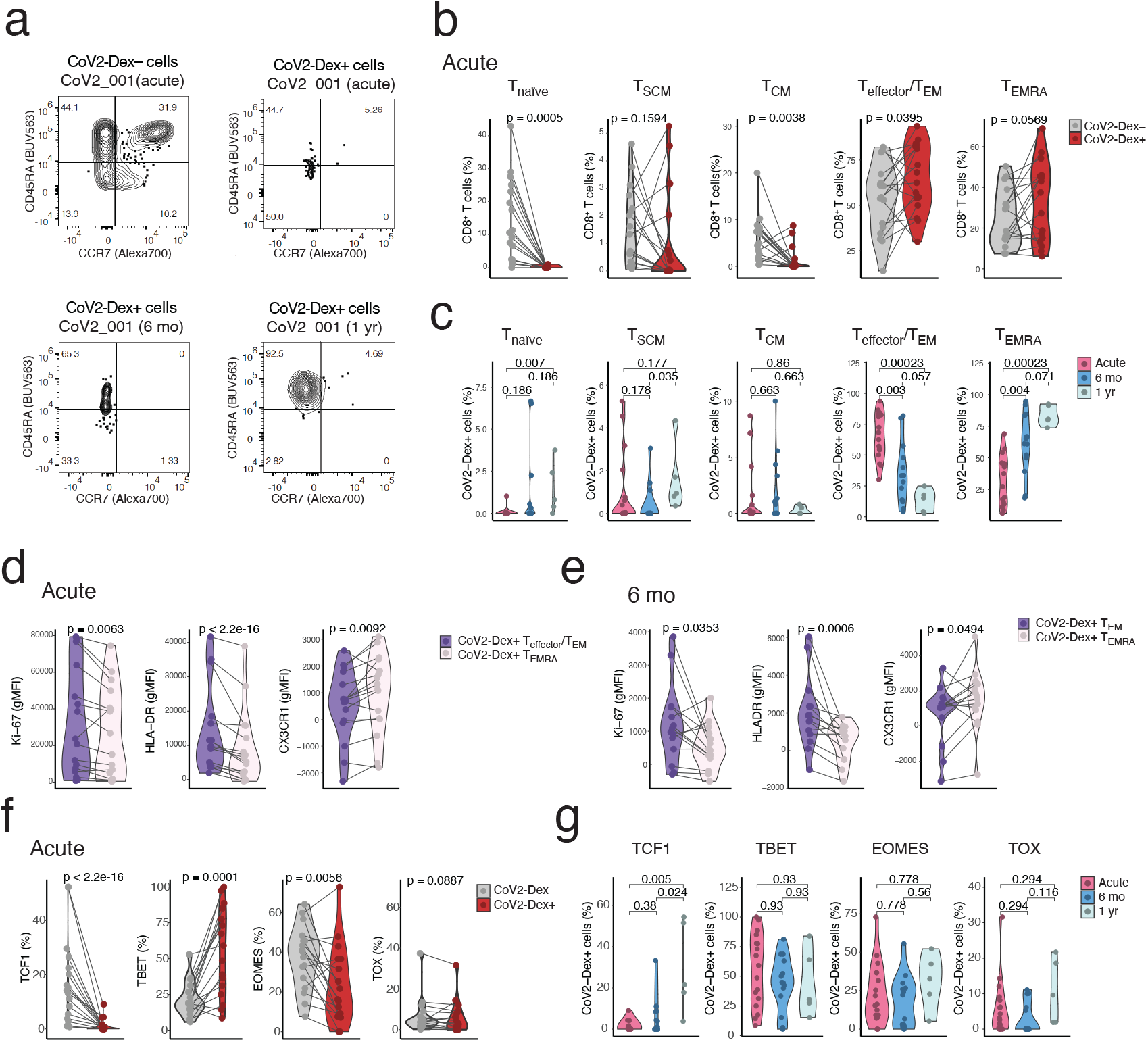
Transition of antigen-specific CD8^+^ T cells to TCF1^+^ CD45RA^+^ effector-memory cells at one year. **a**, Representative plots of CD45RA and CCR7 staining on CoV2-Dex^−^ cells and CoV2-Dex^+^ cells during the acute phase, six months and one year after primary infection. **b**, Percentage of naïve, stem cell memory (T_SCM_), central memory (T_CM_), Effector/Effector memory (T_effector_/ T_EM_) and effector memory T cells re-expressing CD45RA (T_EMRA_) cells among CoV2-Dex^−^ and CoV2-Dex^+^ cells during acute COVID-19. **c**, Percentage of naïve, T_SCM_, T_CM_, T_effector_/ T_EM_ and T_EMRA_ CoV2-Dex^+^ cells in the acute phase as well as six months and one year after infection. **d**, Expression of selected markers on T_effector_/ T_EM_ and T_EMRA_ CoV2-Dex^+^ cells in the acute phase. **e**, Expression of selected markers on T_effector_/ T_EM_ and T_EMRA_ CoV2-Dex^+^ cells six months after primary infection. **f**, Percentage of TCF1^+^, TBET^+^, EOMES^+^ and TOX^+^ cells among CoV2-Dex^−^ and CoV2-Dex^+^ cells during acute COVID-19. **g**, Percentage of TCF1^+^, TBET^+^, EOMES^+^ and TOX^+^ CoV2-Dex^+^ cells in the acute phase as well as six months and one year after infection. **b**,**c**,**d**,**e**,**f**,**g**, Each dot represents independent and unrelated donors. **b**,**d**,**e**,**f**, P-values were calculated with a Wilcoxon signed-rank test. **c**,**g**, P-values were calculated with a Mann-Whitney-Wilcoxon test and corrected for multiple comparisons with the Holm method.

Subsequently, we assessed whether the T_effector_/T_EM_ and T_EMRA_ phenotypes were associated with specific T cell markers, suggesting distinct differentiation states. Indeed, in CoV2-Dex^+^ cells, we observed several differences between the T_effector_/T_EM_ and T_EMRA_ populations (Fig. 3d). CoV2-Dex^+^ T_EM_ cells showed higher Ki-67 and HLA-DR, whereas they had lower abundance of CX3CR1 already during the acute phase. Notably, we observed similar phenotypic changes in CoV2-Dex^+^ T_EM_ cells six months after infection (Fig. 3e).

As T cell phenotypes are driven by specific transcription factors, we assessed the transcription factors TCF1, T-box expressed in T cells (TBET), eomesodermin (EOMES), and thymocyte selection-associated high-mobility group box (TOX), which are known to play important roles in T cell differentiation^26–28^. CoV2-Dex^+^ cells expressed increased TBET in the acute phase (Fig. 3f and Extended Data Fig. 8a), which was maintained for more than a year after infection. Conversely, TCF1 was downregulated during the acute phase and was progressively restored at subsequent timepoints (Fig. 3g and Extended Data Fig. 8b). A difference in TCF1 or TBET expression between specific T_effector_/ T_EM_ and T_EMRA_ was not evident, although specific T_EMRA_ expressed significantly lower EOMES six months after infection (Extended Data Fig. 8c). In summary, our data support a model in which transition to memory is accompanied by progressive CD45RA expression, cessation of rapid proliferation, and acquisition of TCF1^26^.

## Specific signatures identify antigen-specific CD8^+^ T cell memory precursors

To longitudinally track individual antigen-specific T cell clones, we performed TCR sequencing of CoV2-Dex^+^ cells, which revealed several antigen-specific CD8^+^ T cell clones for each epitope investigated (Fig. 4a). Clones were considered antigen-specific if any of the clonal cells were CoV2-Dex^+^ (see Supplementary Dataset 1), and clones that were CoV2-Dex^+^ in the acute phase were considered CoV2-Dex^+^ independently of CoV2-Dex staining at six months after infection, and *vice versa*. As expected, the number of clones detected during convalescence was markedly lower than that detected during the acute phase of infection (Fig. 4a). In most cases, but not all, dominant clones in the acute phase corresponded to the largest clones found in the recovery phase (Fig. 4b). We examined the clones detected in the acute phase that were still present in the convalescent phase (persistent) or became undetectable (non-persistent) (Fig. 4c and Extended Data Fig. 9a). Clone size correlated positively with persistence (Fig. 4d). Interestingly, cells of persistent clones showed a different transcriptional makeup at the acute phase when compared to cells of non-persistent clones (Fig. 4e), which also resulted in a different distribution among the previously identified CD8^+^ T cell clusters (Extended Data Fig. 9b). This effect was robustly seen in different clones and was not due to a few hyper-expanded clones (Extended Data Fig. 9c). Gene set enrichment analysis (GSEA) revealed distinct signatures in persistent vs. non-persistent clones. Persisters were enriched in genes involved in IFN-γ response, IFN-α response, and TNF signaling, whereas non-persisters were enriched in mTOR signaling genes and genes related to mitosis (Fig. 4F). By comparing differentially regulated genes between persistent and non-persistent clones we observed activation (*HLA-DQA1, HLA-DPA1*), terminal differentiation (*KLRG1*) and cytotoxicity (*GZMM, NKG7*) genes to be enriched in persisters, together with CD45RA protein expression determined by TotalSeq™ (Fig. 4g). Conversely, non-persisters showed higher expression of CTLA-4, TIM-3 (*HAVCR2*) and Ki-67 (*MKI67*), as well as the pro-inflammatory cytokine interleukin-32 (Fig. 4g). We also observed differential TCR-Vβ usage between persistent and non-persistent clones (Fig. 4g).

**Figure 4.**
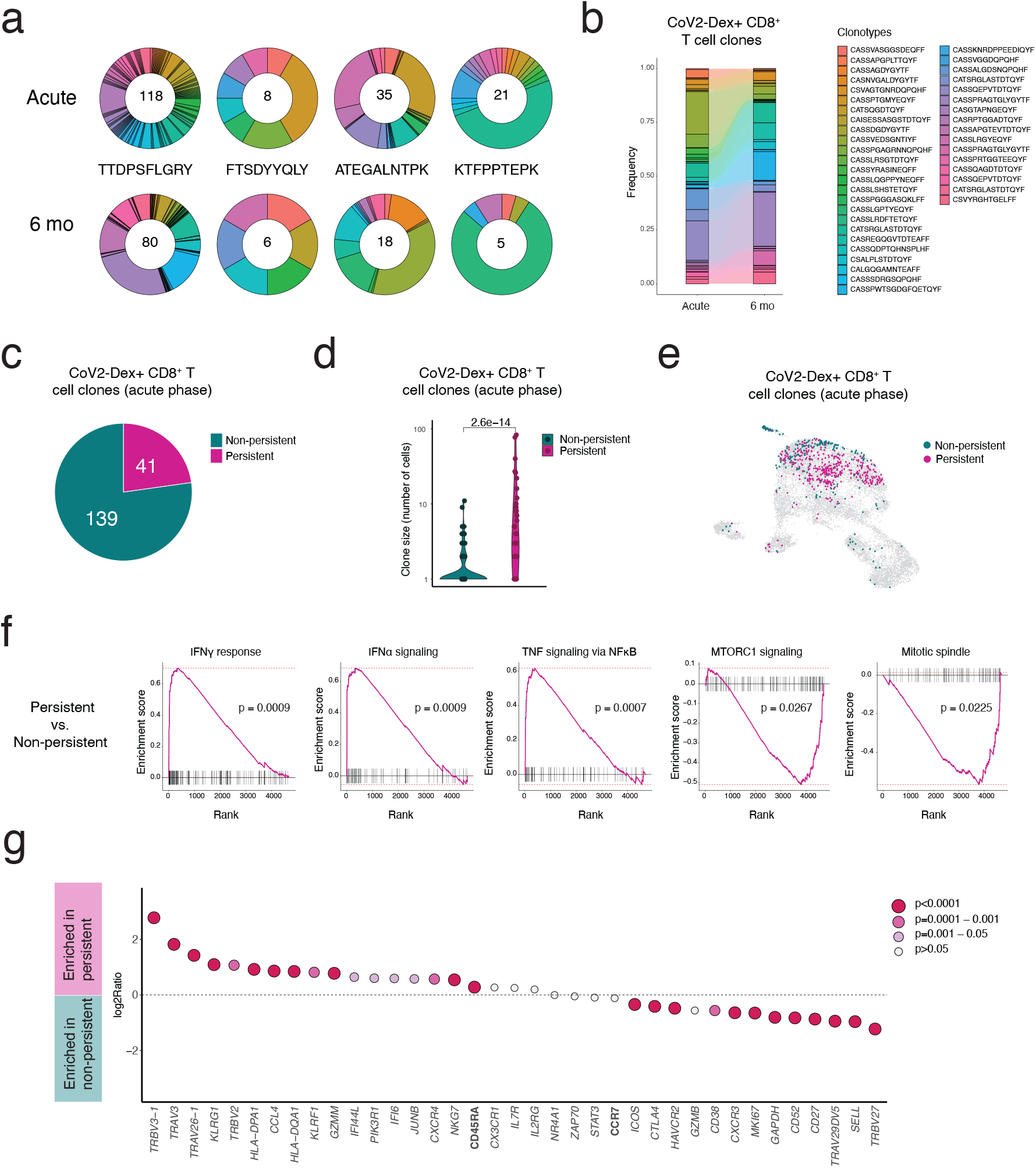
Transcriptional signature of antigen-specific CD8^+^ T cell clones persisting at six months. **a**, Clonotype distribution among CoV2^+^ T cell clones for each of the four epitopes assessed. The number within the circle indicates the number of T cell clones for the given epitope at the given time point. **b**, Alluvial plot showing the relative representation of single clones that were present both during acute COVID-19 and six months after primary infection. **c**, Proportion of clones present during the acute phase that were still detectable six months after primary infection. **d**, Clone size of persisting T cell clones, compared to clones that were not detectable six months after primary infection. The p-value was calculated with a Mann-Whitney-Wilcoxon test. **e**, UMAP plot of T cells belonging to CoV2^+^ T cell clones colored by status (persistent vs. non persistent clones). **f**, Gene set enrichment analysis (GSEA) showing the enrichment of genes associated with cytokine signaling among persistent clones and mTOR signaling as well as proliferation among non-persistent clones. **g**, Expression of selected genes (italics) and CD45RA protein determined by Totalseq™ for persistent vs. non-persistent clones. P-values were adjusted for multiple comparisons.

## Discussion

As hundreds of millions of people worldwide recover from COVID-19, the longevity of functional T cell memory after infection with SARS-CoV-2 will remain a major concern in the coming years. Here, we report for the first time, robust T cell memory one year after infection. Furthermore, this is the first description in humans of a transcriptional and phenotypical trajectory of memory CD8^+^ T cells differentiating from T_effector_/T_EM_ to T_EMRA_ and concomitantly upregulating TCF1 following *de novo* infection with a natural virus. Our data is in agreement with previous reports of memory differentiation in yellow fever virus vaccinees^14,15^, revealing that despite their poor proliferative capacity upon TCR stimulation *in vitro*^29^, CD8^+^ T_EMRA_ cells constitute the main memory phenotype in peripheral blood.

Understanding how the immune system maintains the balance between effector response and memory formation could explain why some infections result in robust and long-lasting T cell memory, whereas others fail to do so. Likely, both pathogen and host factors shape the fate of memory T cells. Our study helps unravel the complexity of these processes by finding a transcriptional signature that correlates with the acquisition of long-lived memory T cells. Somewhat surprisingly, although clone size during the acute infection is generally related to persistence, we find that a strongly proliferative phenotype is associated with clonal contraction and disappearance. Furthermore mTOR signaling, which is likely stimulated by TCR engagement, appears to play a role in instructing the fate of short-lived effector cells by skewing cells away from memory precursors in humans, as previously shown in mice^12^. Conversely, cytokine signaling mark cells destined to become memory, in agreement with previous studies showing the importance of type I IFN signaling for memory generation^30^. Our data suggest formation of memory CD8^+^ T cells to be strongly dependent on a delicate balance between cytokine and TCR signaling during acute infection, which in turn influence outcomes of long-lived memory T cells.

## Methods

### Human subjects and patient characteristics

Following written informed consent, adult patients with symptomatic, RT-qPCR confirmed SARS-CoV-2 infection were recruited in the Canton of Zurich, Switzerland, between April 2 and September 24, 2020. The study was approved by the Cantonal Ethics Committee of Zurich (BASEC 2016-01440). 175 patients donated peripheral blood at the time of inclusion into the study, and 115 and 38 patients donated peripheral blood six months and one year after infection, respectively. Standardized clinical data were collected for all included patients and disease severity was assessed as previously described for this cohort^1–3^. Following HLA class I typing, patients carrying an A*01:01, A*11:01 and/or an A*24:02 allele with sufficient bio-banked samples at two different time-points were selected for the study (*n* = 33). 13 healthy donors carrying A*01:01, A*11:01 and/or an A*24:02 allele were included for comparison.

### Dextramer and Pentamer staining

4*10^6^ peripheral blood mononuclear cells (PBMCs) per patient were first incubated with Human TruStain FcX™ blocking reagent (Biolegend, Cat# 422302) for 10 minutes at 4°C. After washing, cells were incubated with MHC-I dextramers (see Supplementary Table 1 for complete list) in the presence of L-biotin and Herring Sperm DNA according to the manufacturer’s instructions, for 10 minutes at room temperature. Concentrated antibody mixes for either cell sorting or spectral flow cytometry staining were then added to the samples and cells incubated for further 30 minutes at 4°C or 20 minutes at room temperature according to the experiment. For MHC-I pentamer staining, cells were incubated for 10 minutes at 37°C with pentamers (see Extended Table 1 for complete list), a concentrated antibody mix was then added to the samples and cells incubated for further 20 minutes at room temperature. Frozen PBMCs were used throughout the study.

### Spectral Flow Cytometry

After dextramer staining, a concentrated surface staining antibody mix (see Supplementary Table 2 for complete list) was applied without washing and samples were incubated at room temperature for further 20 minutes. After four rounds of washing cells were re-suspended in a fixation permeabilization solution (eBioscience™ Foxp3 / Transcription Factor Staining Buffer) and incubated for 60 minutes at room temperature. After washing an antibody mix for intracellular staining (see Supplementary information for complete list) was added, and cells were incubated for 30 minutes at room temperature. After washing, samples were acquired on a Cytek Aurora spectral flow cytometry.

### Cell sorting

A concentrated antibody mix containing TotalSeq™ antibodies (see Supplementary Table 3 for complete list) was applied after dextramer staining without washing (0.25 μg / 10^6 cells) and cells were incubated at 4°C for 30 minutes. After 4 rounds of washing, cells were re-suspended in PBS with 2% FBS and 2mM EDTA and sorted. For each patient, CoV2-Dex^−^ and CoV2-Dex^+^ cells were sorted approximately in a 10:1 ratio. All CoV2-Dex^+^ cells from each sample were sorted, the corresponding amount of CoV2-Dex_–_ was calculated and sorted in the same tube. Cells from ten patients and the same time point were pooled together, generating four individual sample sets in total: (1) patients CoV2_T001–CoV2_T010, acute; (2) patients CoV2_T001–CoV2_T010, six months after infection; (3) patients CoV2_T011–CoV2_T020, acute; (4) patients CoV2_T011–CoV2_T011-20, six months after infection.

### Additional sample sets (unsorted cells and healthy controls)

We additionally generated two more sample sets: using 5000 unsorted PBMCs from each patient’s sample: (5) patients CoV2_T001–CoV2_T010, six months after infection unsorted; (6) patients CoV2_T011–CoV2_T020, six months after infection unsorted. Finally, using PBMCs from four healthy donors, we generated sample set (7) by sorting and pooling 2000 CD8^+^ T cells from each healthy donor sample.

### scRNAseq library preparation and sequencing

Cells of sample sets 1-7 were analyzed by single cell RNA sequencing utilizing the 5’ Single Cell GEX and VDJ v1.1 platforms (10x Genomics). Each sample set was processed individually. Cell suspensions were pelleted, re-suspended and loaded into the Chromium Chip following the manufacturer’s instructions. 14 cycles of initial cDNA amplification were used for all sets and single cell sequencing libraries for whole transcriptome analysis (GEX), TCR profiling (VDJ), and combined cell surface protein and dCODE Dextramer detection (ADT) were generated. Final libraries were quantified using a Qubit Fluorometer, pooled in a ratio of 5:1:1 (GEX:VDJ:ADT) and sequenced on a NovaSeq 6000 system with following cycle configuration: Read 1: 28bp, Index read 1: 10bp, Read 2: 101bp.

### Single-cell transcriptome analysis

Raw scRNA-seq FASTQ files were aligned to the human GRCh38 genome with Cell Ranger version 5.0.0 with default settings for the ‘cellranger multi’ pipeline (10x Genomics). The reference genome was downloaded from the 10x Genomics website (https://cf.10xgenomics.com/supp/cell-exp/refdata-gex-GRCh38-2020-A.tar.gz) and built as per official release notes (https://support.10xgenomics.com/single-cell-gene-expression/software/release-notes/build#GRCh38_2020A). Every sample set was analyzed with the ‘cellranger multi’ pipeline, which allows to process together the paired GEX, ADT and VDJ libraries for each set. Downstream analysis was conducted in *R* version 4.1.0 with package *Seurat* version 4.0.3 (PMID: 34062119). Cells with < 200 or > 2.500 detected genes and cells with > 10% detected mitochondrial genes were excluded from the analysis.

To be able to investigate possible patient biases, we de-multiplexed cells from patient pools 1-6 based on genetic variants detected within the scRNA-seq reads. For this, we used the tool *souporcell* version 2 (PMID: 32366989). To cluster cells based on their patient specific genetic variants, we merged sample sets 1, 2 and 5 (comprising sorted cells from both time points of patients CoV2_T001–CoV2_T010 and unsorted cells of the same patients) and sets 3, 4 and 6 (comprising cells from both time points of patients CoV2_T011–CoV2_T020 and unsorted cells of the same patients). Then, we executed the souporcell pipeline with option *k=10* (number of clusters to be determined) for each of the two merged sample sets. This analysis allowed us to classify 88% of the cells passing the filtering steps from above into 20 genotype-driven “patient” clusters.

After log normalization and variable feature calculation, independent datasets were integrated using *Seurat’s* anchoring-based integration method. Data scaling, principal component analysis, clustering and UMAP visualizations were performed on the integrated dataset using 15 principal components and a resolution of 0.5 for the shared nearest neighbor clustering algorithm. To define distinct biological features of cell clusters, differential gene expression analyses were done on assay “RNA” of the integrated dataset. *FindAllMarkers* was executed with *logfc*.*threshold* and *min*.*pct* cutoffs set to 0.25. For the analysis of clusters, *FindMarkers* was used with default settings for the comparison of persistent and non-persistent clones. For the differential expression analysis of manually selected genes and cell surface proteins (CD45RA and CCR7), *logfc*.*threshold* and *min*.*pct* cutoffs were set to 0.

For gene set enrichment analysis, the *FindMarkers* function from *Seurat* was first used for the differential expression of genes between cells belonging to persistent and non-persistent clones (using the default Wilcoxon Rank Sum test, with options ‘min.pct=0.1, logfc.threshold = -Inf’, to account also for small expression changes, as long as the genes were expressed in at least 10% of cells of at least one group). The resulting 4701 genes were pre-ranked in decreasing order by the negative logarithm of their p-value, multiplied for the sign of their average log fold change (in R, ‘-log(p_val)*sign(avg_log2FC)’). Gene Set Enrichment Analysis (GSEA)^4^ was performed on this pre-ranked list using the R package *FGSEA* (https://github.com/ctlab/fgsea/)^5^. We used the FGSEA-simple procedure with 100000 permutations and the hallmark gene sets for *Homo sapiens* from the Molecular Signatures Database (https://www.gsea-msigdb.org/gsea/msigdb/index.jsp, made accessible in R by the package *msigdbr*, https://github.com/cran/msigdbr) and set the seed value (‘set.seed(42)’ in R) prior to execution in order to make the results reproducible. The results were filtered for gene sets that were significantly enriched with adjusted p-value < 0.1.

### T cell receptor profiling

Paired chain TCR sequences were obtained through targeted amplification of full-length V(D)J segments during library preparation. Sequence assembly and clonotype calling was done through *cellranger’s* immune profiling pipeline (*cellranger multi*). TCR profiling on filtered contig annotations was done using R package *scRepertoire* version 1.1.4^6^, which assigns TCR nucleotide and amino acid sequences together with clonal frequency counts and a clonotype classification to each cell. Function *combineTCR* was executed with *filterMulti=T* to isolate the top two expressed chains in cell barcodes with multiple chains. Clonotypes were called based on the amino acid sequence of the CDR3 region of TCRα and TCRβ chains. For cells where only one of the two chains could be identified, the available chain was used. Clone calling was done for each sample set independently before integration.

### SARS-CoV-2 peptide loaded dextramer binding of CD8^+^ T cells

To identify SARS-CoV-2-specific CD8^+^ T cells, we used dCODE Dextramers® loaded with viral peptides presented on MHC-I molecules. Initially, two peptides presented on HLA-A*01:01 (FTSDYYQLY from ORF3a and TTDPSFLGRY from ORF1ab), two peptides presented on HLA-A*11:01 (ATEGALNTPK and KTFPPTEP from the nucleocapsid protein) and one peptide presented on HLA-A*24:02 (QYIKWPWYI from the spike protein) were included. To assess unspecific binding, a negative control dextramer (peptide STEGGGLAY presented on HLA-A*01:01) and a general negative control dextramer were included. After analysis of the flow cytometry data, we noticed strong background staining of dextramer A*24:02 (peptide QYIKWPWYI) in samples of healthy donors, indicating unspecific dextramer binding. Thus, we excluded all sequencing counts from this dextramer in the downstream analysis. For other dextramers, cells were considered CoV2-Dex^+^ when the UMI count of a CoV2-Dextramer was >10 and more than five times higher than the UMI count of the negative control in the same cell. Cells that were positive for more than one dextramer according to this classification (< 0.2% of all cells with known TCR) were excluded from the analysis. A TCR clone was considered SARS-CoV-2-specific when at least one cell of the clone was CoV2-Dex^+^.

### Statistics

The Wilcoxon-Mann-Whitney test was used for comparisons of two independent groups. The Wilcoxon signed rank test was used for paired testing. The p-values were adjusted for multiple comparisons with the Holm method. A linear regression model was used to quantify the relationship between variables. Significance was assessed by non-parametric methods unless otherwise specified. All tests were performed two sided. Analyses were performed with R (version 4.0.0 or 4.1.0).

## Data availability

The sequencing dataset generated during the current study have been deposited at <u>zenodo.org</u> and are available at https://doi.org/10.5281/zenodo.5119633. Flow cytometry datasets are available from the corresponding author on reasonable request.

## Code availability

The code generated during the current study is available at https://github.com/TheMoorLab.

## Acknowledgments

We thank Sara Hasler for her assistance with patient recruitment and coordination, Esther Baechli, Alain Rudiger, Melina Stüssi-Helbling, Lars C. Huber, and Dominik J. Schaer for their support in patient recruitment, and the members of the Boyman Laboratory for helpful discussions. Graphical representations were generated with BioRender.com.

## Author contributions

S.A. designed and performed the experiments, analyzed data and contributed to data interpretation. J.M. performed the scRNAseq experiments, analyzed data and contributed to data interpretation. Y.Z. contributed to experiments, patient recruitment and data collection. C.C. and P.T. contributed to patient recruitment and data collection.

M.E.R contributed to patient recruitment and clinical management. S.B.S. contributed to the analysis of scRNAseq data. J.N. contributed to study design and patient recruitment. A.E.M. contributed to experiment design and data interpretation. O.B. conceived the project, contributed to experimental design and data interpretation. S.A. and O.B. wrote the manuscript with contribution by J.M. and S.B.S. All authors edited and approved the final draft of the article.

## Additional information

**Supplementary Information** is available for this paper.

Correspondence and requests for materials should be addressed to O.B.

## References

1. Crotty, S. & Ahmed, R. Immunological memory in humans. Semin. Immunol. 16, 197–203 (2004).

2. Sallusto, F., Lanzavecchia, A., Araki, K. & Ahmed, R. From vaccines to memory and back. Immunity 33, 451–463 (2010).

3. Saad-roy, C. M. et al. Immune life history, vaccination, and the dynamics of SARS-CoV-2 over the next 5 years. Science (80-.). 818, 811–818 (2020).

4. Jalkanen, P. et al. COVID-19 mRNA vaccine induced antibody responses against three SARS-CoV-2 variants. Nat. Commun. 12, 1–11 (2021).

5. Chemaitelly, H. et al. mRNA-1273 COVID-19 vaccine effectiveness against the B.1.1.7 and B.1.351 variants and severe COVID-19 disease in Qatar. Nat. Med. (2021) doi:10.1038/s41591-021-01446-y.

6. Sette, A. & Crotty, S. Adaptive immunity to SARS-CoV-2 and COVID-19. Cell 184, 861–880 (2021).

7. Tan, A. T. et al. Early induction of functional SARS-CoV-2-specific T cells associates with rapid viral clearance and mild disease in COVID-19 patients. Cell Rep. 34, 108728 (2021).

8. Szabo, P. A. et al. Longitudinal profiling of respiratory and systemic immune responses reveals myeloid cell-driven lung inflammation in severe COVID-19. Immunity 54, 797–814 (2021).

9. Dan, J. M. et al. Immunological memory to SARS-CoV-2 assessed for up to 8 months after infection. Science (80-.). 371, 6529 (2021).

10. Zuo, J. et al. Robust SARS-CoV-2-specific T cell immunity is maintained at 6 months following primary infection. Nat. Immunol. 22, 620–626 (2021).

11. Bonifacius, A. et al. COVID-19 immune signatures reveal stable antiviral T cell function despite declining humoral responses. Immunity 54, 340–354 (2021).

12. Araki, K. et al. mTOR regulates memory CD8 T-cell differentiation. Nature 460, 108–112 (2009).

13. Langenkamp, A. et al. T cell priming by dendritic cells: Thresholds for proliferation, differentiation and death and intraclonal functional diversification. Eur. J. Immunol. 32, 2046–2054 (2002).

14. Akondy, R. S. et al. The Yellow Fever Virus Vaccine Induces a Broad and Polyfunctional Human Memory CD8 + T Cell Response. J. Immunol. 183, 7919–7930 (2009).

15. Akondy, R. S. et al. Origin and differentiation of human memory CD8 T cells after vaccination. Nature 552, 362–367 (2017).

16. Saini, S. K. et al. SARS-CoV-2 genome-wide T cell epitope mapping reveals immunodominance and substantial CD8+ T cell activation in COVID-19 patients. Sci. Immunol. 6, 1–23 (2021).

17. Peng, Y. et al. Broad and strong memory CD4+ and CD8+ T cells induced by SARS-CoV-2 in UK convalescent individuals following COVID-19. Nat. Immunol. 21, 1336–1345 (2020).

18. Weiskopf, D. et al. Phenotype and kinetics of SARS-CoV-2-specific T cells in COVID-19 patients with acute respiratory distress syndrome. Sci. Immunol. 5, eabd2071 (2020).

19. Grifoni, A. et al. Targets of T Cell Responses to SARS-CoV-2 Coronavirus in Humans with COVID-19 Disease and Unexposed Individuals. Cell 181, 1489-1501.e15 (2020).

20. Braun, J. et al. SARS-CoV-2-reactive T cells in healthy donors and patients with COVID-19. Nature 587, 270–274 (2020).

21. Le Bert, N. et al. SARS-CoV-2-specific T cell immunity in cases of COVID-19 and SARS, and uninfected controls. Nature 584, 457–462 (2020).

22. Le Bert, N. et al. Highly functional virus-specific cellular immune response in asymptomatic SARS-CoV-2 infection. J. Exp. Med. 218, e20202617 (2021).

23. Hou, S., Hyland, L., Ryan, K. W., Portner, A. & Doherty, P. C. Virus-specific CD8+ T-cell memory determined by clonal burst size. Nature 369, 652–654 (1994).

24. Surh, C. D. & Sprent, J. Homeostasis of Naive and Memory T Cells. Immunity 29, 848–62 (2008).

25. Raeber, M. E., Zurbuchen, Y., Impellizzieri, D. & Boyman, O. The role of cytokines in T-cell memory in health and disease. Immunol. Rev. 283, 176–193 (2018).

26. Escobar, G., Mangani, D. & Anderson, A. C. T cell factor 1: A master regulator of the T cell response in disease. Sci. Immunol. 5, eabb9726 (2020).

27. Intlekofer, A. M. et al. Effector and memory CD8+ T cell fate coupled by T-bet and eomesodermin. Nat. Immunol. 6, 1236–1244 (2005).

28. Khan, O. et al. TOX transcriptionally and epigenetically programs CD8(+) T cell exhaustion. Nature 571, 211–218 (2019).

29. Geginat, J. et al. Proliferation and differentiation potential of human CD8. Blood 101, 4260–4266 (2003).

30. Kolumam, G. A., Thomas, S., Thompson, L. J., Sprent, J. & Murali-Krishna, K. Type I interferons act directly on CD8 T cells to allow clonal expansion and memory formation in response to viral infection. J. Exp. Med. 202, 637–650 (2005).

## References

1. Cervia, C. et al. Systemic and mucosal antibody responses specific to SARS-CoV-2 during mild versus severe COVID-19. J. Allergy Clin. Immunol. 147, 545-557.e9 (2020).

2. Chevrier, S. et al. A distinct innate immune signature marks progression from mild to severe COVID-19. Cell Reports Med. 2, 100166 (2021).

3. Adamo, S. et al. Profound dysregulation of T cell homeostasisand function in patients with severe COVID-19. Allergy 1–16 (2021) doi:10.1111/all.14866.

4. Subramanian, A. et al. Gene set enrichment analysis: A knowledge-based approach for interpreting genome-wide expression profiles. Proc. Natl. Acad. Sci. U. S. A. 102, 15545–15550 (2005).

5. Korotkevich, G. et al. Fast gene set enrichment analysis. (2016) doi:10.1101/060012.

6. Borcherding, N., Bormann, N. L. & Kraus, G. scRepertoire: An R-based toolkit for single-cell immune receptor analysis. F1000Research 9, 1–17 (2020).

