## Extended Data Figures 1-9 for "CD8^+^ T cell signature in acute SARS-CoV-2 infection identifies memory precursors"

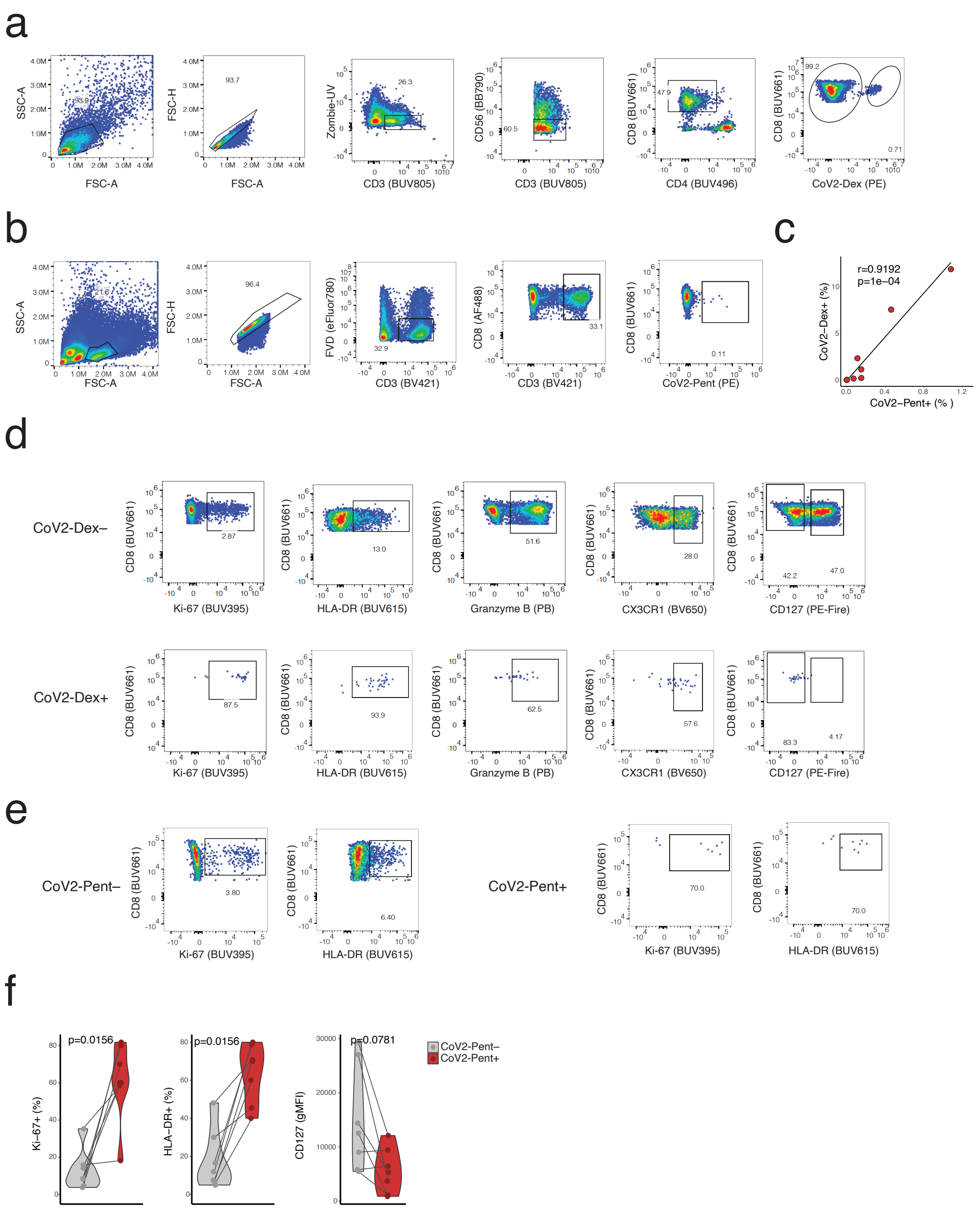


**Extended Data Fig.1. Gating strategy antigen specific CD8⁺ T cells and MHC-I pentamer staining. a**, Gating strategy CoV2-Dex⁺ cells. **b**, Gating strategy CoV2-Pent⁺ cells. **c**, Linear regression of frequency of CoV2-Dex⁺ cells as a function of frequency of CoV2-Pent⁺ cells. **d**, Gating strategy phenotypical analysis of CoV2-Dex⁺ compared to CoV2-Dex⁻ cells. **e**, Gating strategy phenotypical analysis of CoV2-Pent⁺ compared to CoV2-Pent⁻ cells. **f**, Frequency of Ki-67+, HLA-DR+ and CD127 levels cells among CoV2-Pent$^-$ (grey) and CoV2-Pent$^+$ (red). P-values were calculated with a Wilcoxon signed-rank test.

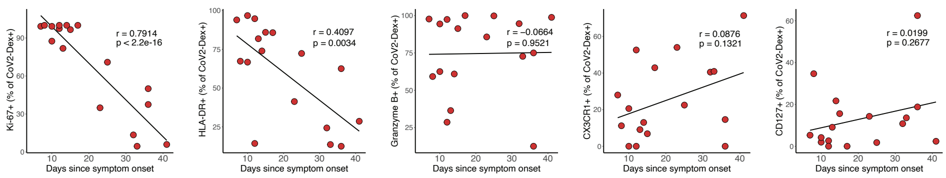


**Extended Data Fig. 2**. **Pseudolongitudinal course of key markers on antigen-specific CD8$^+$ T cells.** Linear regression of frequency of Ki-67$^+$, HLA-DR$^+$, Granzyme B$^+$, CX3CR1$^+$ and CD127$^+$ cells among CoV2-Dex$^+$ cells as a function of time since symptom onset in the acute phase.

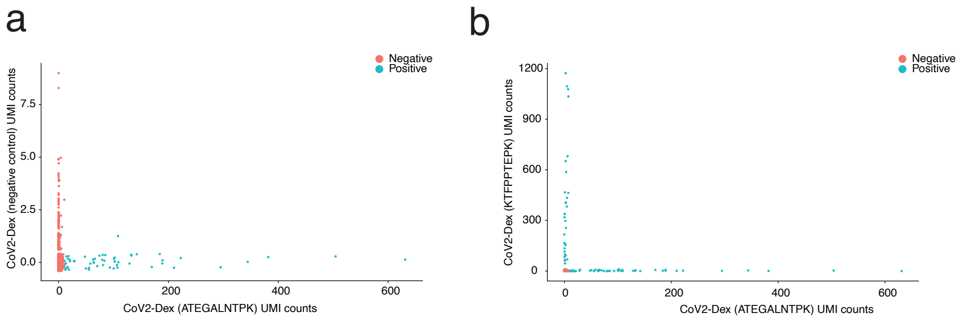


**Extended Data Fig. 3. Definition of CoV2-Dex+ cells. a,** Unique molecular identifier (UMI) counts for CoV2-Dex HLA-A*11:01 (ATEGALNTPK) vs. UMI counts for negative control dextramer; cells defined as CoV2-Dex HLA-A*11:01 (ATEGALNTPK)$^{+}$ are depicted in blue. **b**, (UMI) counts for CoV2-Dex HLA-A*11:01 (ATEGALNTPK) vs. UMI counts for CoV2-Dex HLA-A*11:01 (KTFPPTEPK).

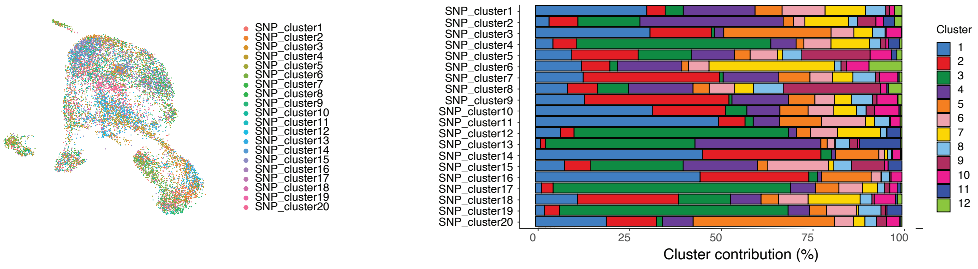


**Extended Data Fig. 4. Single patient contribution to individual clusters**. Uniform manifold approximation and projection (UMAP) plot colored by patient ID (left) and cluster distribution for single patients (right).

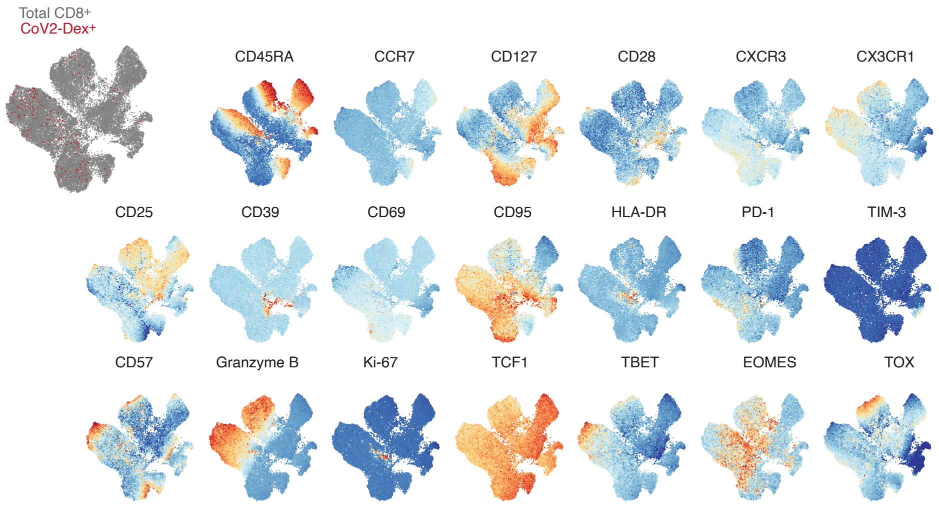


**Extended Data Fig. 5. Phenotype of antigen-specific CD8$^+$ T cells six months after infection.** UMAP plots of marker expression for up to 2000 CD8$^+$ T cells from each sample analyzed by spectral flow cytometry. Regions with high expression of specific markers appear red. Overlay of CoV2-Dex$^+$ cells (red) and total CD8$^+$ T cells (grey) is shown on the upper left.

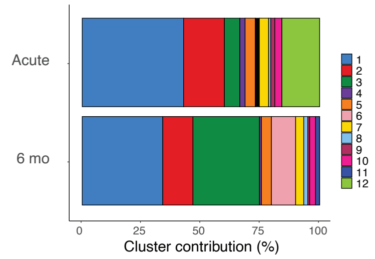


**Extended Data Fig. 6. Cluster composition of CoV2-Dex$^{+}$ cells during acute and recovery phase.** Cluster composition of CoV2-Dex$^{+}$ CD8$^{+}$ T cells in the acute phase vs. six months after infection.

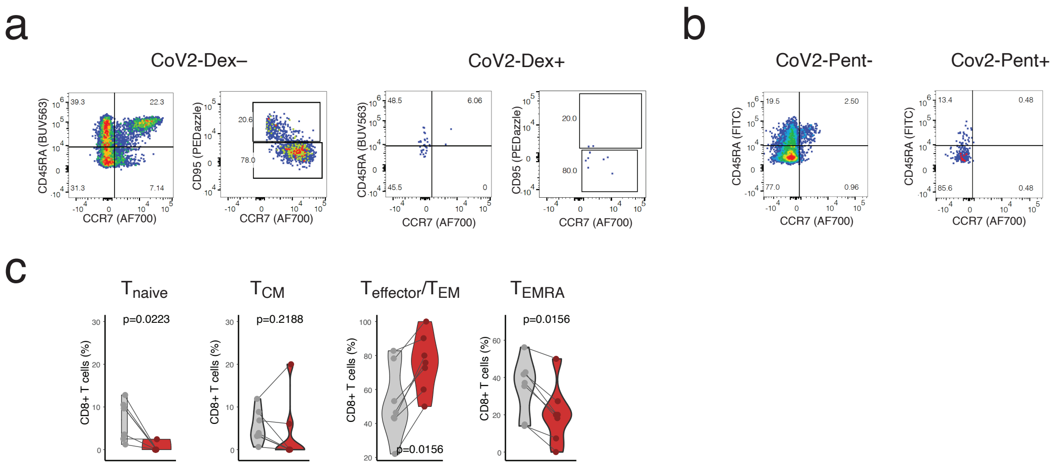


**Extended Data Fig. 7. Memory phenotypes in CoV2-Dex$^+$ and CoV2-Pent$^+$ cells. a,** Gating strategy naïve, stem cell memory ($T_{SCM}$), central memory ($T_{CM}$), Effector/Effector memory ($T_{effector}$/ $T_{EM}$) and effector memory T cells re-expressing CD45RA ($T_{EMRA}$) cells among CoV2-Dex$^-$ and CoV2-Dex$^+$ cells. **b**, Gating strategy naïve, central memory ($T_{CM}$), Effector/Effector memory ($T_{effector}$/ $T_{EM}$) and effector memory T cells re-expressing CD45RA ($T_{EMRA}$) cells among CoV2-Pent$^-$ and CoV2-Pent$^+$ cells. **c**, Percentage of naïve, $T_{CM}$, $T_{effector}$/ $T_{EM}$ and $T_{EMRA}$ cells among CoV2-Pent$^-$ (grey) and CoV2-Pent$^+$ (red) cells during acute COVID-19.

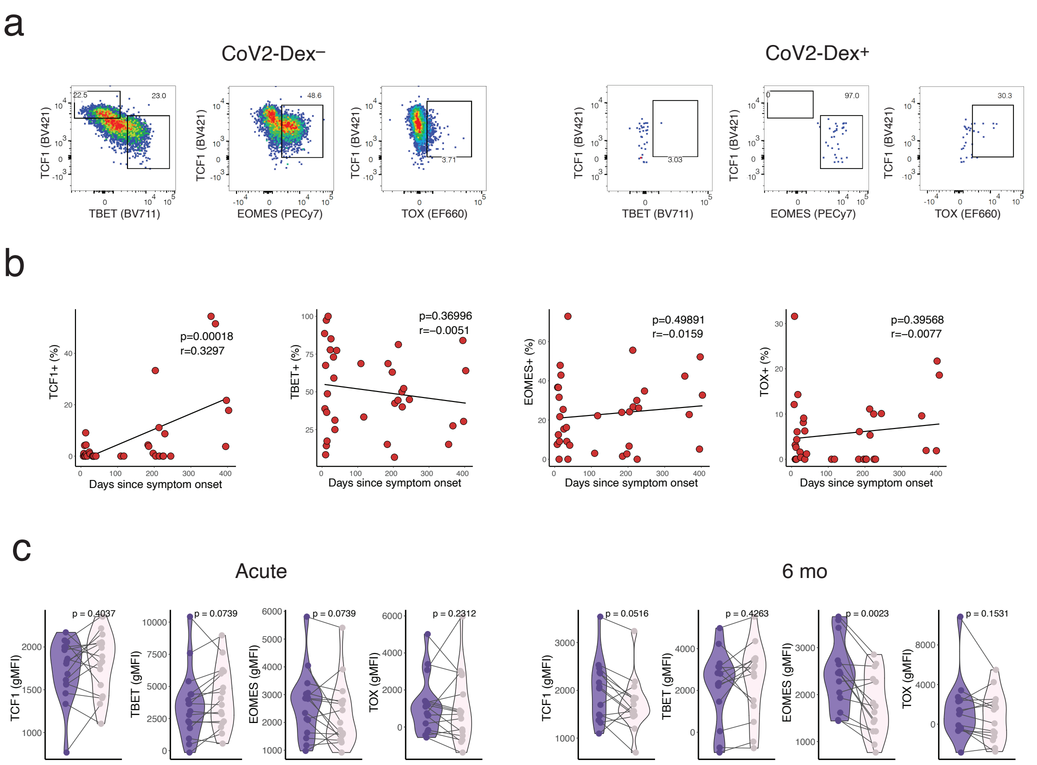


**Extended Data Fig. 8. Gating strategy and pseudo-longitudinal time course of transcription factors. a**, Gating strategy TCF1$^+$, TBET$^+$, EOMES$^+$, TOX$^+$ cells among CoV2-Dex$^-$ and CoV2-Dex$^+$ cells. **b**, Linear regression of frequency of strategy TCF1$^+$, TBET$^+$, EOMES$^+$, TOX$^+$ cells among CoV2-Dex$^+$ cells as a function of time since symptom onset. **c**, Expression of transcription factors on $T_{effector}$/ $T_{EM}$ and $T_{EMRA}$ CoV2-Dex$^+$ cells in the acute phase and six months after infection.

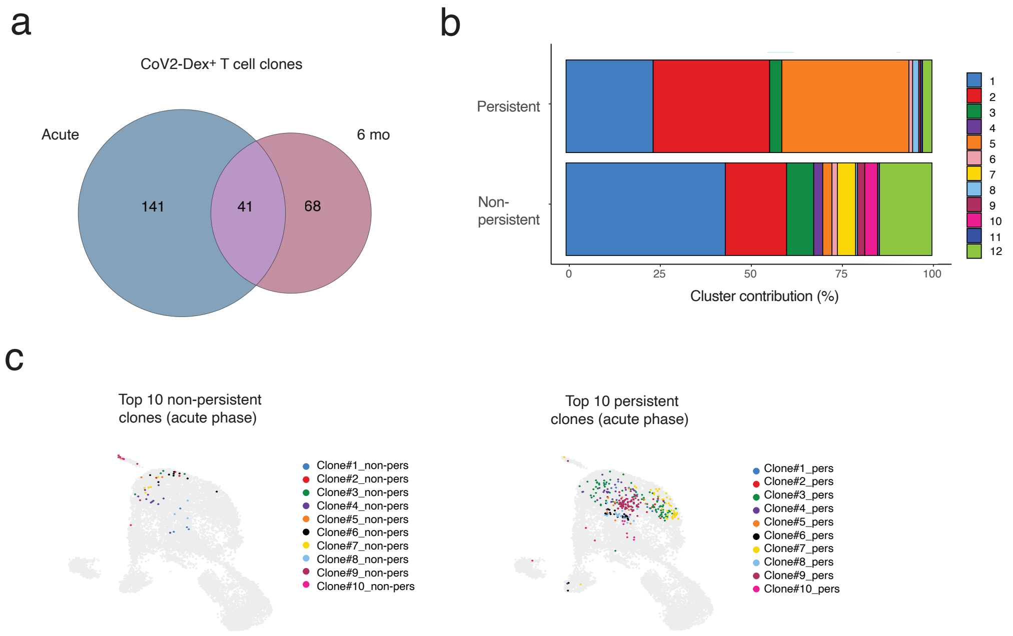


**Extended Data Fig. 9. Clonality and phenotype of persistent vs. non-persistent CoV2-Dex$^{+}$ T cell clones. a**, Venn diagram showing overlapping clones in the acute phase and six months after infection. **b**, Cluster composition of persistent vs. non-persistent CD8$^{+}$ T cell clones. **c**, UMAP plots showing top five persistent CoV2-Dex$^{+}$ CD8$^{+}$ T cell clones and top five non-persistent CoV2-Dex$^{+}$ CD8$^{+}$ T cell clones.
