## Supplementary Tables 1-3 for "CD8^+^ T cell signature in acute SARS-CoV-2 infection identifies memory precursors"

Supplementary Table 1: SARS-CoV2-directed Dextramers and Pentamers

| **Type** | **HLA allele** | **Peptide** | **Fluorophore** | **Nucleotide Tag** | **Manufacturer** | **Cat. #** |
| --- | --- | --- | --- | --- | --- | --- |
| Pent | A01:01 | FTSDYYQLY | PE | - | ProImmune | F4355-2A-E |
| Pent | A11:01 | KTFPPTEPK | PE | - | ProImmune | F4356B-2A-E |
| Dextr | A01:01 | FTSDYYQLY | PE | AGTCGCGCAGTCTGG | Immudex | WA5973-PfBC0602 |
| Dextr | A01:01 | TTDPSFLGRY | PE | TCTCTAGGTGAGTGG | Immudex | WA5974-PfBC0603 |
| Dextr | A11:01 | ATEGALNTPK | PE | CAAGAACCACGCCGT | Immudex | WD5981-PfBC0605 |
| Dextr | A11:01 | KTFPPTEPK | PE | CGGAATCACAATCCG | Immudex | WD5981-PfBC0606 |
| Dextr | A24:02 | QYIKWPWYI | PE | AGGCGCCGTGGTGTT | Immudex | WF5952-PfBC0607 |
| Dextr | A01:01 | STEGGGLAY | PE | ATACTTGTCATCTAT | Immudex | WA3579-PfBC0619 |
| Dextr | Genral | - | PE | CCTGCACGGCCAAGC | Immudex | Ni3233- PfBC0618 |

Supplementary Table 2: Spectral flow cytometry staining antibodies

| **Cell marker** | **Fluorophore** | **Manufacturer** | **Cat. #** |
| --- | --- | --- | --- |
| Ki-67 | BUV 395 | BD | 564071 |
| Zombie UV | UV450 | Biolegend | 423107 |
| CD4 | BUV496 | BD | 564652 |
| CD45RA | BUV563 | BD | 565703 |
| HLA-DR | BUV615 | BD | 751142 |
| CD8 | BUV661 | BD | 741683 |
| CD28 | BUV737 | BD | 564438 |
| CD3 | BUV 805 | BD | 612893 |
| TCF-7/TCF-1 | BV421 | BD | 566692 |
| Granzyme B | Pacific blue | Biolegend | 515408 |
| CXCR3 | BV510 | Biolegend | 353725 |
| PD-1 | BV605 | Biolegend | 329923 |
| CX3CR1 | BV650 | Biolegend | 341625 |
| TBET | BV711 | Biolegend | 644819 |
| CD39 | BV785 | Biolegend | 328239 |
| CD69 | FITC | Biolegend | 310904 |
| CD57 | PerCP Cy5.5 | Biolegend | 359621 |
| CD56 | BB790-P | BD | 624296 (custom) |
| CD95 | PE Dazzle | BD | 562395 |
| CD25 | PECy5 | Biolegend | 302608 |
| CD127 | PE-Fire 700 | Biolegend | 351365 |
| EOMES | PECy7 | Invitrogen | 25-4877-41 |
| TOX | EF 660 | Invitrogen | 50-6502-80 |
| CCR7 | AF700 | BD | 561143 |
| TIM-3 | APC fire | Biolegend | 345043 |

Supplementary Table 3: Cell sorting staining antibodies

| **Cell marker** | **Fluorophore** | **Manufacturer** | **Cat. #** |
| --- | --- | --- | --- |
| CD4 | Pacific Blue | Biolegend | 344620 |
| CD56 | BV510 | Biolegend | 318339 |
| CD3 | BV785 | Biolegend | 300472 |
| CD8 | AF488 | Biolegend | 344716 |
| CD39 | APC | Biolegend | 328209 |
| Fixable viability dye | EF 780 | Invitrogen | 1 65-0865-14 |
| CD45RA | TotalSeq^TM^ | Biolegend | 304163 |
| CCR7 | TotalSeq^TM^ | Biolegend | 353251 |
